# Evolutionarily distinct dengue virus type 2 with a unique type-specific neutralization determinant

**DOI:** 10.64898/2026.11.10.765652

**Authors:** Sandra V. Mayer, Evandro R. Winkelmann, Ana Coello Escoto, Jack Dorman, Michael K. McCracken, Douglas Fritz, Ralph Baric, Longping V. Tse, Richard Herbert, Derek A.T. Cummings, Stephen S. Whitehead, Sonja M. Best, Patrick Dolan, John Aaskov, Gregory D. Gromowski, Leah C. Katzelnick

## Abstract

Dengue viruses 1-4 (DENV1-4) mostly circulate in endemic cycles involving humans and *Aedes* mosquitoes. However, rare dengue outbreaks originate from non-human primates. These spillovers are caused by sylvatic virus genotypes that are closely related to endemic strains of the same serotype, but some are caused by strains that fall between the four canonical serotypes on phylogenetic trees. It is not clear whether these genotypic differences are reflected in corresponding antigenic differences. Here, we characterized the antigenic features of QML-22, a divergent strain most closely related to DENV2. On an antigenic map, QML-22 clustered with other globally representative DENV2 strains. QML-22 was not as well neutralized by antisera raised against endemic DENV2 and was better neutralized by antisera from DENV1, 3, and 4 infections than other strains of DENV2. QML-22 had a cluster of distinct amino acid residues in domain I of the envelope (E) protein, with only scattered point mutations elsewhere in the envelope. A potently neutralizing anti-DENV2 antibody that targets this site in domain I of the E protein, 3F9, had >1000-fold lower neutralization of QML-22 than other endemic DENV2 strains, whereas other cross-reactive and DENV2-specific antibodies neutralized QML-22 and endemic strains to similar titers. When antisera from non-human primates infected with QML-22 or DENV2 reference strain S16803 were depleted with the heterologous virions, homologous neutralization was unaffected but heterologous neutralization was reduced, suggesting homologous neutralization was due to distinct epitopes. Evolutionary analyses showed the 3F9 epitope was under relaxed selection pressure compared to other neutralizing epitopes and that QML-22 had unique amino acids in this epitope relative to the likely common ancestor of all DENV2. These findings provide insight into the antigenic diversification of DENV and the possible evolution of new serotypes.

**Author summary:** Dengue viruses that are most distantly related to the four canonical serotypes originated in Borneo, suggesting this island is a possible site of early dengue virus diversification as well as continued evolution within an unidentified host species. Here, we find divergent strain QML-22 from Borneo shares many neutralizing epitopes with DENV2 but has a distinct epitope in an important region affecting potent DENV2 neutralization. This region of the virus also is variable within DENV2 and in other DENV serotypes, highlighting an important site of dengue virus diversification and evolution. The spillover threat of viruses like QML-22 remains unknown, but ecological changes in Borneo may increase this risk.

## Introduction

Among the arthropod-borne viruses, the four dengue viruses (DENV1-4) represent the greatest threat to human health. DENV is transmitted endemically by *Aedes* mosquitoes and half of the world population is at risk of infection, mostly in tropical and sub-tropical areas [1]. Collectively, DENV1-4 cause an estimated 50 million cases of dengue annually and ∼500,000 severe infections, characterized by vascular leakage, hemorrhage, shock, and organ failure [2]. DENV1-4 are positive-sense, single-stranded RNA viruses encoding three structural proteins, the capsid, envelope (E), and pre-membrane protein (prM), and seven non-structural proteins [3]. Each virion contains 180 E monomers arranged as dimers in an icosahedral virion, and E is the main target of protective and pathogenic antibodies.

Early challenge and serological studies grouped DENV into four distinct serotypes [4] but later analyses revealed multiple genotypes within each serotype that can differ by 6-10% at the nucleotide level [5]. Recent work has further divided the genotypes into major and minor lineages, some of which circulate globally and others that are regionally restricted [6]. Antigenic diversity exists among strains of a given serotype [7–10], and while not thought to be common, homotypic reinfection resulting in disease can occur, especially against a dissimilar genotype of the same serotype [11–13]. The possibility of homotypic reinfection may be greater following vaccination, as vaccines can induce a more antigenically restricted response than natural infection [14]. In a phase 3 clinical trial, a dengue vaccine elicited protection only against the DENV4 genotype used in the vaccine [15, 16]. Analyses using antigenic cartography have also shown that some endemic and sylvatic DENV1-4 are highly antigenically similar to other serotypes, indicating increased susceptibility to cross-reactive antibodies [17]. Thus, antigenic and phylogenic distance are only loosely correlated, with distinct strains potentially more antigenically similar than phylogenetic distance would suggest [17]. DENV1-4 can also evolve to become more cross-reactive, corresponding to larger epidemics [18], potentially because specific antigenic distances between sequential infecting viruses increases disease risk [19]. Binding of cross reactive antibodies, but without potent neutralization, can augment disease severity by increasing infection of Fc*γ* receptor-bearing myeloid cells, a mechanism known as antibody-dependent enhancement [20]. Notably, mutations in the E and prM protein can increase sensitivity to cross-reactive, enhancing antibodies, especially those targeting the fusion loop, by reducing virus maturation (cleavage of the prM protein) as well as increasing viral dynamics (the degree to which the virion samples different conformational states) [21–23].

The four DENV serotypes are thought to have diverged genetically more than 1,000 years ago, moved from sylvatic cycles into those involving humans in the last few hundred years, and further diversified into genotypes and lineages in the last century [5]. DENV has been identified in sylvatic cycles in Southeast Asia and West Africa, which may represent lineages that continued to circulate in non-human primates (NHPs) following the emergence of each serotype into endemic cycles in humans [24]. Yet, because sylvatic DENV2 in Asia and Africa form a phylogenetic cluster distinct from endemic strains, it has also been proposed that sylvatic DENV2 was introduced into African NHPs from Asia by human movement [24]. In addition, representatives of highly divergent lineages of DENV have been isolated in humans, which fall between the four canonical DENV serotypes [25–29].

Remarkably, four highly divergent viruses were recovered from people who had been in rural settings on the island of Borneo, suggesting spillover from sylvatic cycles of transmission. Brun2014 was isolated in Australia from a traveler who had visited Brunei in 2014 and was classified as antigenically most like DENV1 based testing with virus-specific antibodies [25]. However, it constituted the most genetically distinct DENV1, with only 83% nucleotide identity with other DENV1 strains and an estimated common ancestor ∼315 years ago, whereas all other endemic and sylvatic DENV1 strains share an ancestor ∼110 years ago [25]. DKE-121, isolated from a farmer in Sarawak, Malaysia in 2007 [26], is even more phylogenetically distinct, and has up to 12% amino acid divergence in the E protein compared with endemic DENV4 viruses. Neutralization and animal challenge experiments in mice and NHPs showed antisera from DKE-121 infected animals only partly neutralized and protected against endemic and sylvatic DENV4 viruses; further, antisera from DENV4 infected animals had limited neutralization of DKE-121. However, immunization of NHPs with DENV4 prevented DKE-121 infection and tetravalent human vaccine antisera also neutralized DKE-121 [26], suggesting DENV4 immunity in NHPs and humans is protective against DKE-121.

Two other divergent strains, QML-22 and the recently identified UDS37/9, are most similar to DENV2. UDS37/9 was isolated from a patient with severe dengue in Sarawak, Malaysia in 2024 [27, 28]. QML-22 was isolated from an Australian traveler who visited Sabah, Malaysia on the island of Borneo in 2015 and developed dengue-like symptoms [29]. UDS37/9 is more similar to the endemic and sylvatic DENV2 genotypes [27], whereas QML-22 is as phylogenetically distinct from other DENV2 strains as DKE-121 from DENV4 [29]. However, it is not known whether the genetic differences in QML-22 result in distinct antigenic properties. In the present study, we use genetic and evolutionary analyses, antigenic cartography, monoclonal antibody neutralization, and antibody depletion experiments to characterize QML-22 and compare it to endemic and sylvatic DENV1-4 strains.

## Results

### QML-22 is phylogenetically distinct but shares antigenic features with other DENV2 strains

To place QML-22 in the context of currently observed DENV sequence diversity, we made a phylogeny of QML-22 and a globally representative sample of DENV1-4 full protein-encoding nucleotide sequences (**Fig. 1A**). Like DKE-121, the divergent DENV4 strain, QML-22 forms a phylogenetic intermediate, more different than a canonical genotype but less different than the serotypes. At the amino acid level, the E protein of QML-22 had an average of 10.7% divergence compared to sylvatic DENV2 strains and 9.6% relative to endemic DENV2 strains, but this is still less than the minimum difference between serotypes (22.4%) (**Fig. 1B**). Although the E protein of QML-22 shared more similarity with endemic strains as compared to other sylvatic strains, the full polyprotein QML-22 was slightly more different from the endemic strains (14.2%) than the sylvatic strains (14.0%).

**Figure 1.**
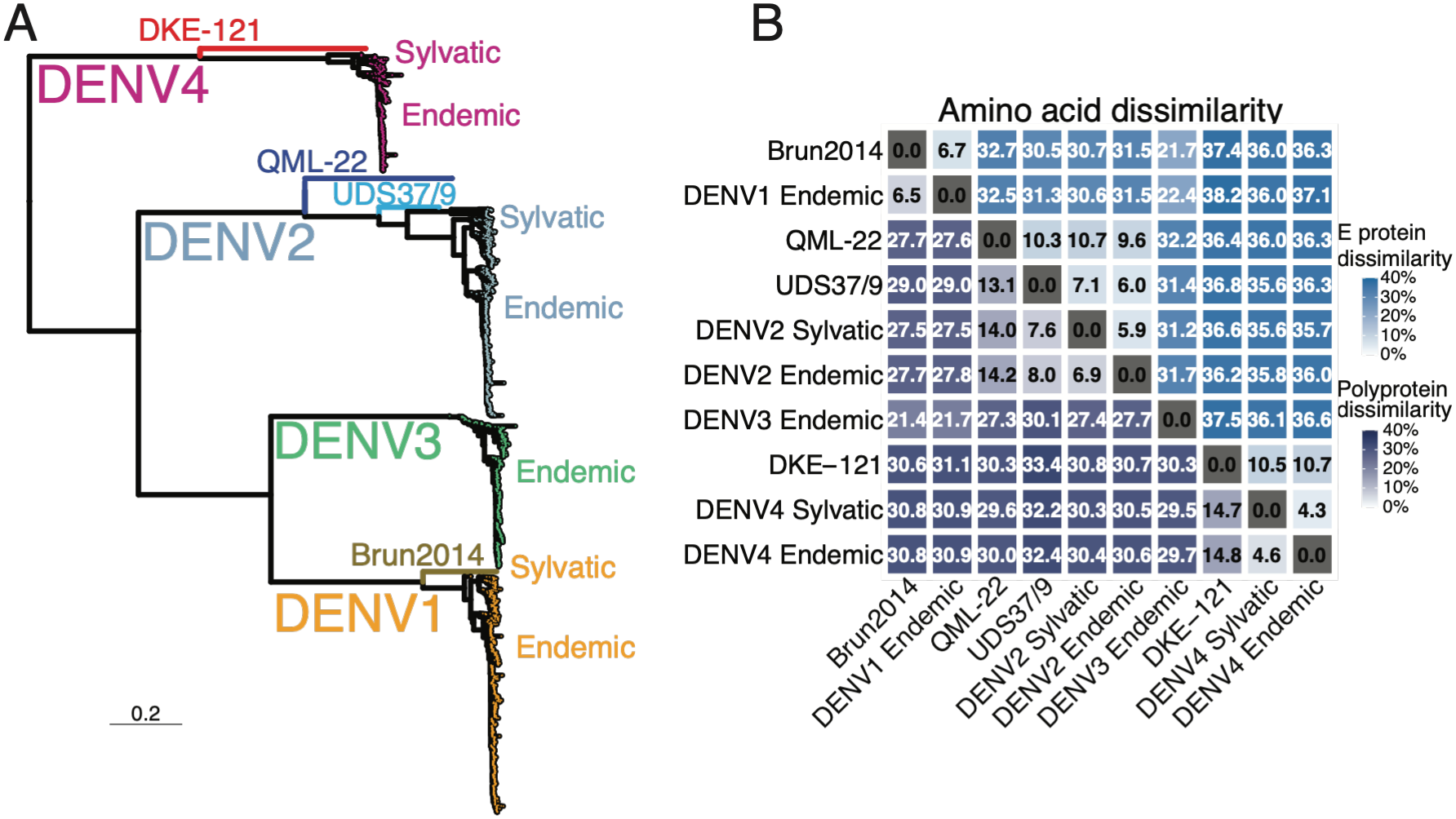
Phylogenetic and genetic relatedness of QML-22 to sylvatic and endemic DENV1-4. (**A**) Maximum likelihood phylogenetic tree of the protein-encoding nucleotide sequences for QML-22, other divergent strains, and globally representative endemic and sylvatic DENV1-4 strains (n=2,969). The tree was estimated in IQ-Tree, with GTR+F+R7 providing the best fit. (**B**) Percent dissimilarity in the amino acid sequences of the envelope proteins (lighter blue shades, upper triangle) or full polyproteins (navy shades, bottom triangle) of known divergent strain and the averages for endemic and sylvatic DENV1-4.

We then used antigenic cartography to define the antigenic relatedness of QML-22 to other endemic and sylvatic DENV1-4 strains that have been characterized previously [17, 18]. Two rhesus macaques were inoculated with QML-22 to produce homologous antisera, which was used for evaluation of neutralization of QML-22 and representative DENV1-4. On an antigenic map of globally representative strains [17], QML-22 clustered with other DENV2 strains. However, QML-22 was located close to the map center, indicating it was better neutralized by antisera raised against other serotypes than most other DENV2 strains (**Fig. 2A**). When QML-22 was added to a larger map including n=365 additional contemporary endemic strains mostly from Thailand [18], QML-22 was also among the top 10% most cross-neutralized, i.e., antigenically central, DENV2 strains (**Fig. 2B**).

**Figure 2.**
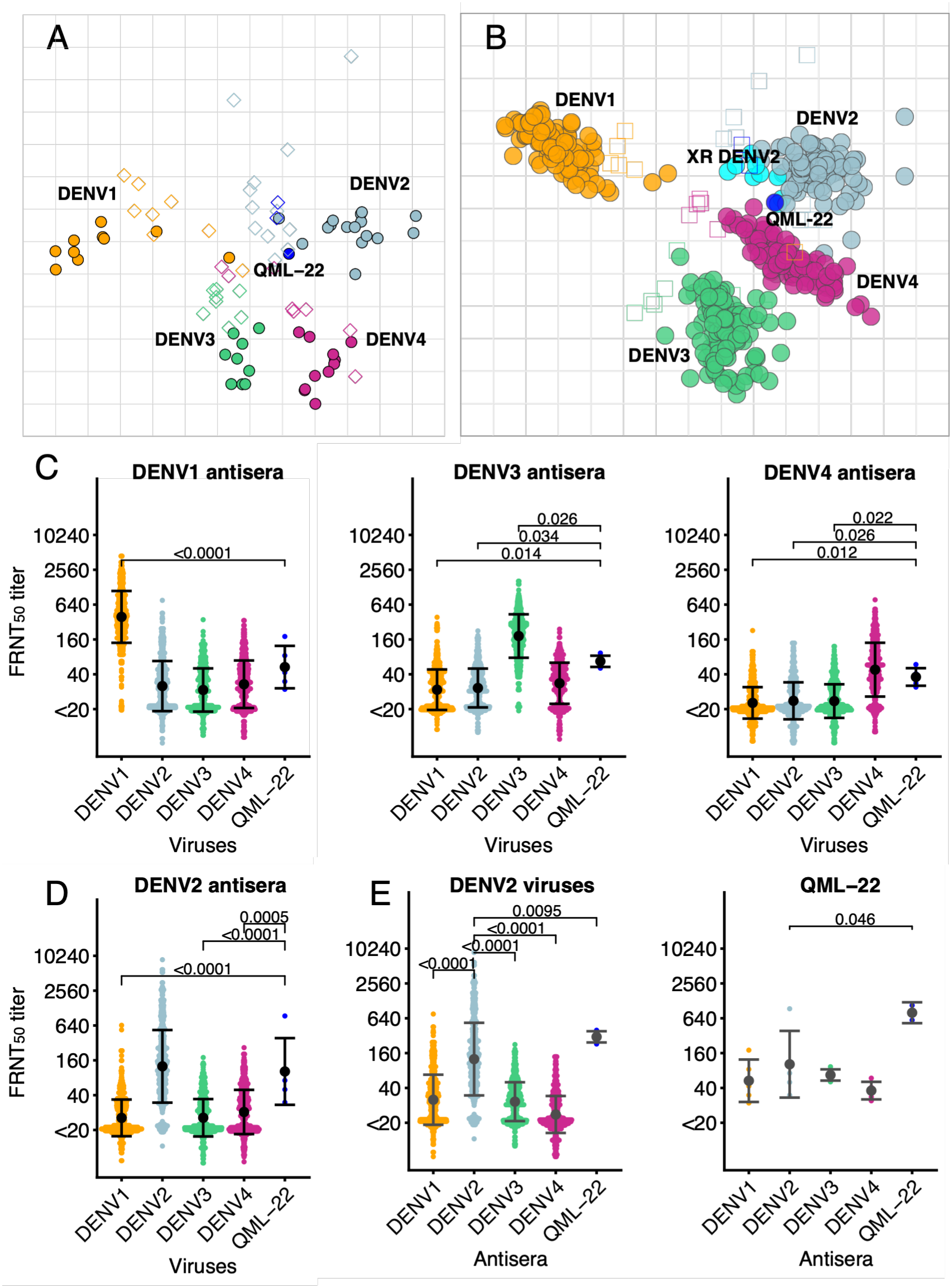
Antigenic relationships between QML-22 and endemic and sylvatic DENV1-4. Antigenic maps of (**A**) n=47 diverse DENV1-4 viruses [17] or (**B**) n=412 DENV1-4 with oversampling of strains from Thailand [18], and QML-22. Viruses were tested for neutralization by antisera produced by infecting NHPs with diverse DENV strains (n=38 antisera), including QML-22. Colored circles show positions of viruses and squares show the positions of antisera. The top 10% most cross-reactive DENV2 strains are colored cyan and defined by distance from the antigenic map center in 3 dimensions [18]. QML-22 is among those top 10%, but colored dark blue. Each grid-square corresponds to a two-fold dilution in the FRNT, which was performed on C6/36 cells. The neutralizing antibody titers (n= 8,681) used for making the antigenic maps were plotted separately for antisera from primary NHP infection with DENV1, DENV3, or DENV4 (**C**) or DENV2 (**D**). The x-axis indicates the serotype of the strains tested, with QML-22 shown separately. (**E**) The same titers were also plotted to visualize how DENV2 viruses or QML-22 were neutralized across the antisera. Each neutralization titer is shown as an individual point, and error bars reflect means +/- 1 standard deviation. Differences in neutralization for each plot were evaluated with a mixed-effects model to account for repeat measures, using the Kenward-Roger method to estimate the degrees of freedom and Tukey method to adjust for multiple comparisons.

Interrogation of the neutralizing antibody titers used for making the maps confirmed these patterns. Antisera from DENV3 and DENV4 infected NHPs neutralized QML-22 to a higher titer than other strains of heterotypic serotypes (**Fig. 2C**). Yet, antisera from DENV2 infected NHPs still neutralized QML-22 to the same degree as other DENV2 viruses (**Fig. 2D**). Visualized a different way, antisera to DENV2 had significantly higher neutralization titers against DENV2 than antisera to DENV1, 3, and 4, whereas antisera raised to DENV1-4 had relatively balanced neutralization of QML-22 (**Fig. 2E**). The exception was QML-22 sera, which strongly neutralized both QML-22 and other DENV2 strains.

Our initial hypotheses for why QML-22 was similarly neutralized by antisera to all four serotypes were that (1) QML-22 was an intrinsically more dynamic virus, resulting in increased susceptibility to low-quality cross-reactive antibodies such as those targeting the fusion loop, (2) QML-22 better presented ancestral cross-reactive epitopes such as those targeted by envelope dimer epitope (EDE) antibodies, and (3) QML-22 lost, or never developed, a serotype-specific epitope. To test these hypotheses, we compared neutralization of QML-22 to neutralization of endemic DENV2 strains that fell near the antigenic map center (Tonga) or periphery (Cambodia). To test if QML-22 had greater intrinsic dynamics, we measured how fusion loop antibodies neutralized each virus when incubated together for either 1.5 or 4 hours before inoculation of cells, with larger increases with time indicating greater virus dynamics. The fusion loop antibodies similarly neutralized all strains, with modest and variable fold-change in neutralization with time, suggesting the differences between strains in virion dynamics were minimal (**Fig. 3A**).

**Figure 3.**
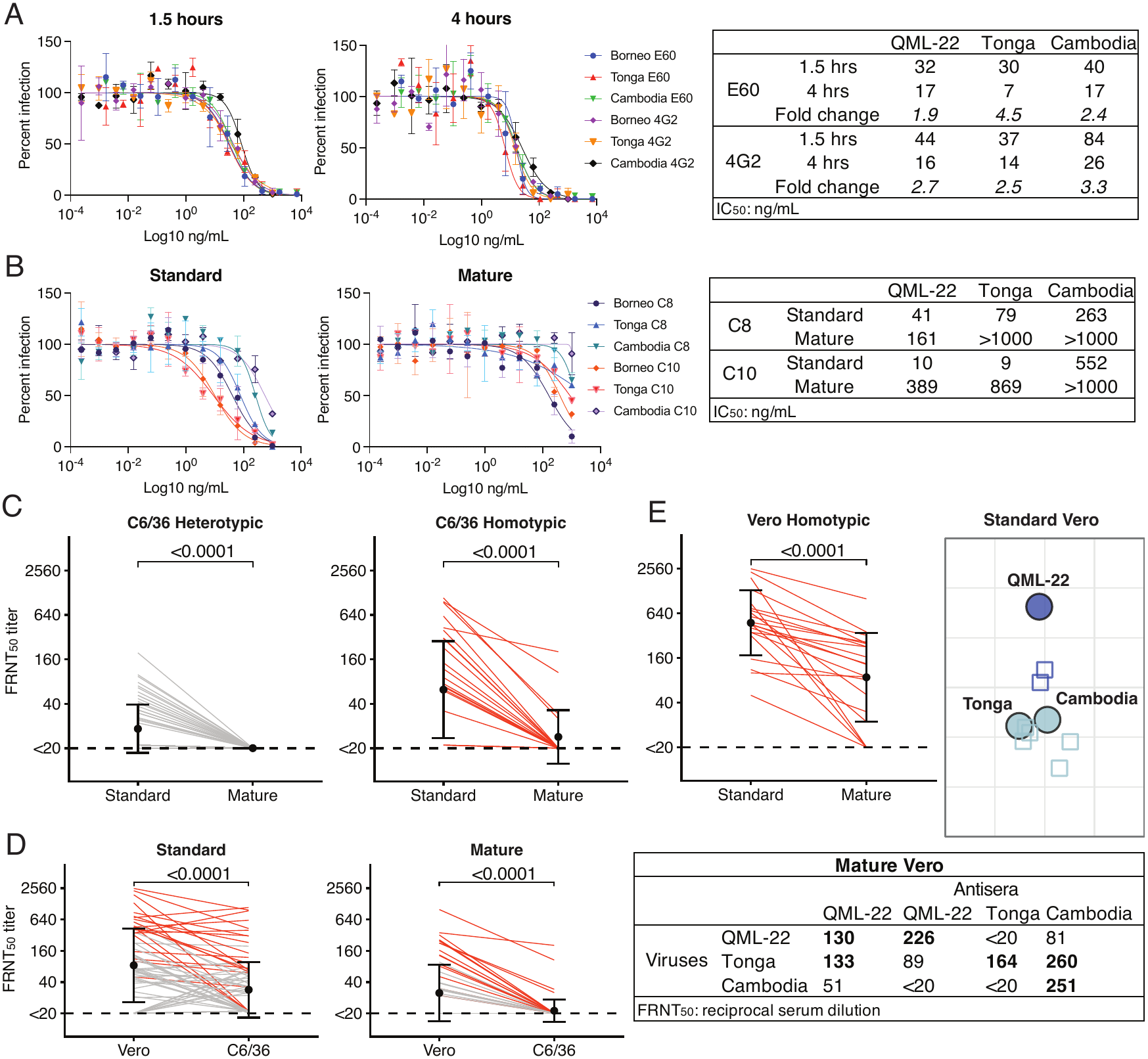
Evaluation of differences in virion structural dynamics and susceptibility to cross-reactive and type-specific antibodies for QML-22 and two endemic DENV2 strains, Tonga and Cambodia. Neutralization of (**A**) the DENV2 strains by fusion loop antibodies E60 and 4G2 when incubated for 1.5 or 4 hours prior to inoculation of cells, or (**B**) partially immature (standard) and mature stocks of each DENV2 strain by EDE antibodies (C8 and C10). Plots show neutralization curves with means +/- 1 standard deviation for technical duplicates. Tables show the concentration (ng/mL) at which 50% neutralization was observed relative to the negative control, with fold-change shown for the fusion loop antibody experiment. Neutralization of (**C**) mature and standard DENV2 strains by homotypic and heterotypic NHP antisera on C6/36 cells, (**D**) mature and standard DENV2 strains on Vero versus C6/36 cells, and (**E**) standard and mature DENV2 strains by homotypic antisera on Vero cells. Lines show paired antiserum-virus titrations between conditions (red, homotypic; grey, heterotypic), and error bars reflect means +/- 1 standard deviation. Differences in neutralization for each plot were evaluated with a mixed-effects model to account for repeat measures, using the Kenward-Roger method to estimate the degrees of freedom and Tukey method to adjust for multiple comparisons. Panel E also shows an antigenic map made using the DENV2 antisera against standard DENV2 strains, and a table of the matched DENV2 antisera titrated against mature DENV2 strains. Each grid-square on the antigenic map corresponds to a 2-fold difference in the neutralization assay. Viruses are labeled, and sera are colored by infecting virus (endemic DENV2 versus QML-22).

To determine if QML-22 may have better retained conserved cross-reactive epitopes, we tested how the potently neutralizing EDE antibodies neutralized the strains. EDE antibodies C8 and C10 better neutralized the QML-22 and Tonga strains than the Cambodia strain (**Fig. 3B**). To further test if QML-22 was more sensitive to potent cross-reactive antibodies, we performed the same experiments using mature stocks of each strain grown in a furin over-expressing cell line. The EDE antibodies had lower but still potent titers against mature QML-22 but could not neutralize the mature Cambodia strain and had lower neutralization of the mature Tonga strain. Together, these results suggest QML-22 was more efficiently neutralized by potent EDE antibodies.

Finally, to evaluate whether QML-22 might also have distinct strain-specific epitopes, we produced mature stocks of each strain and titrated them against the panel of NHP antisera. However, neutralization of the mature viruses was so low that some antisera could not even neutralize the homotypic serotype (**Fig. 3C**). Previous studies of virus maturation have used mammalian cells and have shown Vero cells are mostly infected by mature virus [30]; we thus repeated our evaluation of the viruses by performing the titrations on Vero cells. Most antisera had higher neutralization titers against standard as well as mature viruses on Vero compared to C6/36 cells (**Fig. 3D**). We thus used the Vero titers to evaluate how antisera against DENV2 and QML-22 neutralized the three DENV2 strains (**Fig. 3E**). On an antigenic map made with the standard Vero titers, QML-22 clustered separately from the Tonga and Cambodia strains, indicating it was less efficiently neutralized by the endemic DENV2 antisera. In the mature Vero neutralization data, the antisera raised to Cambodia and Tonga had lower neutralization to QML-22, whereas the antisera raised against QML-22 had high neutralization of the QML-22 strain, with some neutralization also observed to the Tonga strain. This observation indicated differential neutralization of QML-22 by antisera raised to endemic DENV2 strains and that Vero neutralization assays may provide greater insight into epitope-specific differences between strains.

### QML-22 has unique amino acids associated with limited neutralization by a potent antibody

To identify potentially distinct residues in QML-22, we evaluated the amino acid differences between the E proteins of QML-22 other globally representative DENV2 strains. QML-22 was different from all other DENV2 strains at 24 positions, and 98% different at 12 additional positions (**Table S1**). While unique amino acids were observed across the E protein, many mutations were observed in E domain I. These mutations clustered in the center of the epitope for 3F9, a potently neutralizing human anti-DENV2 monoclonal antibody (**Fig. 4A**). Prior work showed that 3F9 lost neutralization of a reference DENV2 strain when the residues at positions 160, 161, 162, 163, 170, 171, 173, 174, 176, 177, and 180 in the E protein were changed to the corresponding amino acids in DENV4 [31]. Among these 11 amino acids, QML-22 was different from all other DENV2 strains at three sites (E161D, T176E, and T180S) and over 95% different at four additional sites (K160Q, I162V, A173T, E174A). The non-conservative amino acid changes, including K160Q, A173T, E174A, and T176E, clustered together on the surface of domain I. QML-22 also differed from over 95% of DENV2 strains at I139V, V140A, V181I, T182S. While these are conservative changes, they are directly beneath, or adjacent to, the 3F9 epitope, suggesting they may also contribute to the structure or function of the epitope.

**Figure 4.**
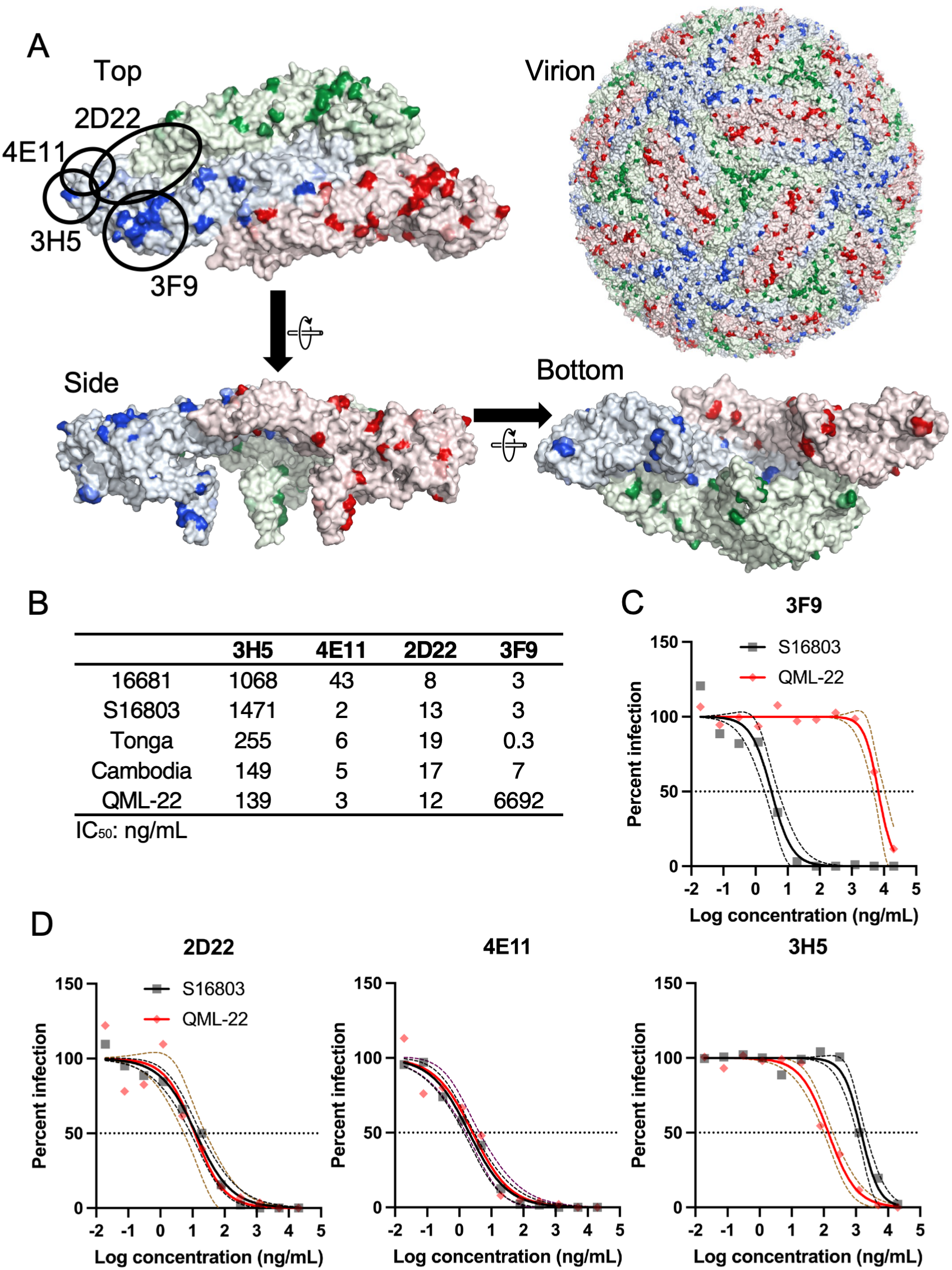
Mapping of a unique QML-22 neutralizing epitope. (**A**) Top, side, and bottom view of the E protein dimer (Protein Data Bank #: 6ZQU, mature DENV2 virus at 3.1 angstrom resolution) as well as the full virion. Monomers are colored by chain in the raft (blue, green, and red for the chains in the 5, 3, or 2-fold vertices, respectively). The approximate locations of each known monoclonal antibody epitope are circled in black and labeled. Residues are shaded by how distinct QML-22 is from other strains (n=843), with the most saturated color indicating close to 100% difference and lighter shading indicating difference from a smaller percentage of DENV2 strains. (**B**) Neutralization (IC_50_, expressed in ng/mL) of QML-22 and diverse DENV2 strains by a panel of DENV2-specific neutralizing antibodies targeting diverse epitopes measured by FRNT on Vero cells. Percent infection of S16803 versus QML-22 when titrated against serial dilutions of monoclonal antibodies (**C**) 3F9, and (**D**) 2D22, 4E11, and 3H5. Curves show estimates and dotted lines indicate 95% confidence intervals, based on technical duplicates for each dilution.

To test the potential effect of these differences in E domain I and other sites in the E protein, we used 3F9 and other DENV2-neutralizing monoclonal antibodies with defined epitopes and assessed their ability to neutralize the QML-22, Tonga, and Cambodia strains as well as DENV2 reference viruses 16681 and S16803 on Vero cells by FRNT (**Fig. 4B**). The 3F9 antibody potently neutralized all strains except QML-22, to which it had titers 1,000 to more than 20,000-fold lower than to the other DENV2 strains (**Fig. 4C**). Antibodies 3H5 and 4E11, which bind distinct but overlapping epitopes on E domain III [32, 33], neutralized QML-22 similarly to or better than the other strains (**Fig. 4D**). 2D22, which binds parts of E domain I and III on one monomer and E domain II of a neighboring monomer within an E dimer [34], also neutralized all strains similarly. These observations point to the 3F9 epitope as one of the neutralizing determinants that distinguished QML-22 from S16803 and other endemic DENV2 strains.

### QML-22 has both virus-specific and shared DENV2-specific neutralizing epitopes

A prior study by Gallichotte et al. demonstrated that for many individuals, the greatest proportion of DENV2-specific neutralization could be explained by antibodies targeting E domain III, and thus more like the 2D22 epitope, than E domain I, more like the 3F9 epitope [31]. However, in some individuals, a large proportion of DENV2 neutralization could be explained just by E domain I, with an average of 54% in natural immune antisera and 30% in vaccine antisera [31]. Given that QML-22 had distinct amino acids that affected recognition by the 3F9 antibody, and our mature neutralization data suggesting QML-22 may be differentially neutralized by antisera raised to endemic DENV2 as compared to QML-22, we evaluated whether infection with QML-22 versus a DENV2 reference strain resulted in distinct neutralization profiles using antibody depletion experiments. Antisera from NHPs experimentally infected with QML-22 or DENV2 reference strain S16803 [35] were depleted with QML-22, S16803, and a mixture of DENV1, DENV3, and DENV4. The responses of depleted antisera were compared to mock-depleted and un-depleted antisera from the S16803 and QML-22 inoculated NHPs, which had high neutralizing antibody titers to both DENV2 strains but mostly undetectable titers to DENV1, DENV3, and DENV4 (**Table S2**). Following depletion, antisera from a flavivirus-naïve NHP had background levels of IgG binding for all ELISA antigens tested (**Fig. S1**). Matched antigen depletion and ELISA combinations, e.g., depletion with S16803 and testing by ELISA against S16803, demonstrated substantial removal of binding antibodies, whereas antisera depleted with bovine serum albumin (BSA), a negative control, retained DENV IgG binding capacity. Mismatched antigen depletion and ELISA combinations, e.g., depletion with QML-22 tested by ELISA against S16803, removed variable levels of DENV IgG.

Antisera that had been depleted with each antigen were then tested for neutralizing antibody titers to S16803 and QML-22 by FRNT on Vero cells (**Fig. 5**). Antisera depleted with BSA had similar neutralizing antibody titers to un-depleted antisera across conditions. Depletion with DENV1, DENV3, and DENV4 had no effect on homotypic or heterotypic DENV2 neutralization for any antiserum tested. Only the homotypic depletion antigen substantially reduced neutralizing antibody titers (Fig. 5A and **B**). Specifically, antisera from S16803-inoculated animals depleted with S16803 had reduced neutralization of S16803 (18.2 and 31.0-fold) but depletion with QML-22 yielded neutralization of S16803 that was equivalent with BSA or DENV1, DENV3, and DENV4 depletion. Similarly, antisera from QML-22 inoculated animals depleted with QML-22 had reduced neutralization of QML-22 (4.8 and 6.8-fold) compared smaller changes in QML-22 neutralization following depletion with S16803 (2.4 and 3.0-fold). The limited effect of depletion with the mismatched DENV2 strain, S16803 or QML-22, on homotypic neutralization indicated that S16803 and QML-22 had distinct strain-specific neutralization determinants.

**Figure 5.**
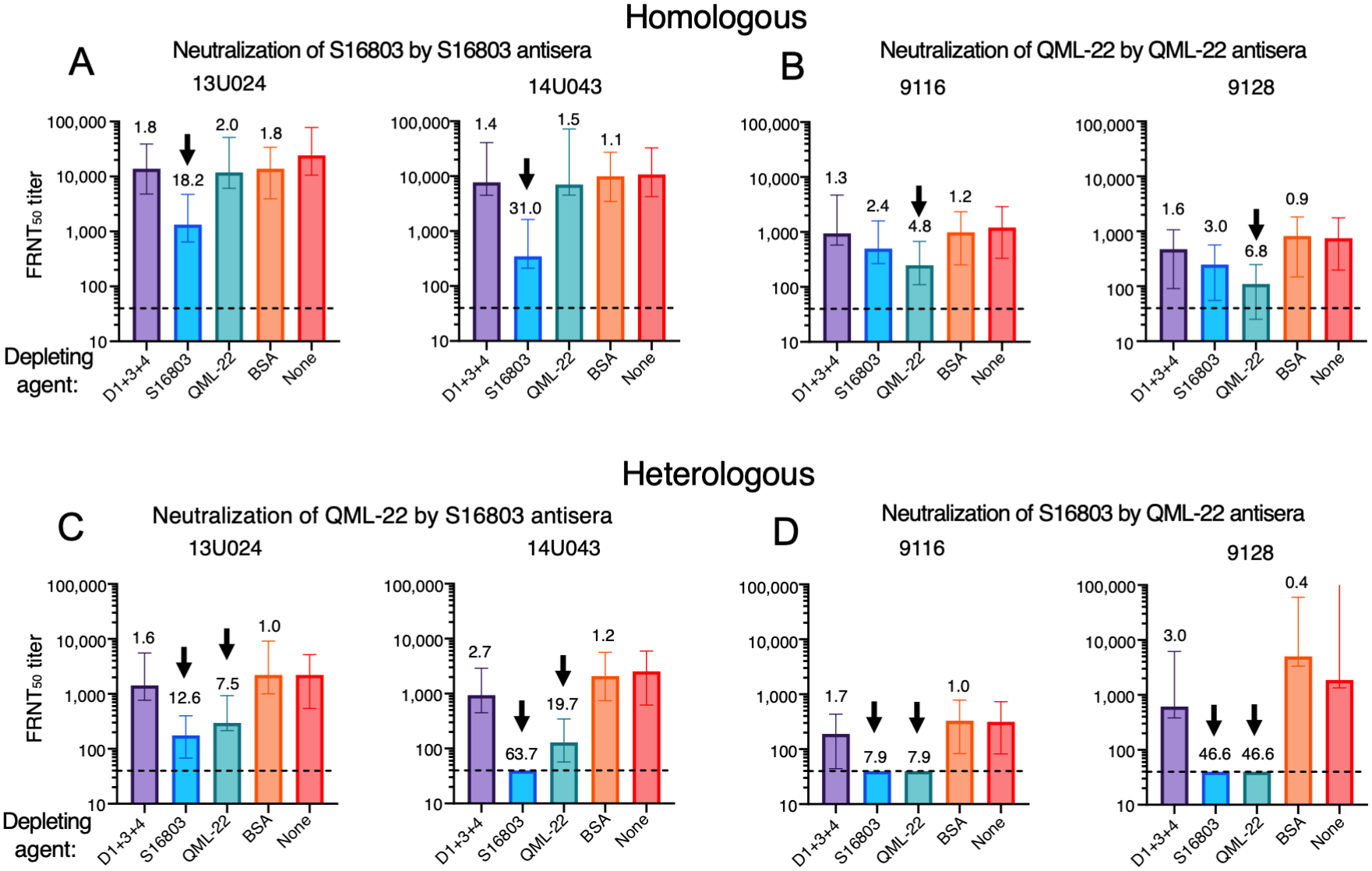
Homologous and heterologous DENV2 neutralization of QML-22 and S16803 under various antibody-depletion conditions. The FRNT_50_ measured on Vero cells for each antibody-depleted antiserum (animal named above each plot) and controls. The depletion conditions are indicated below the bars for each graph, and include depletion with whole virus of DENV1, DENV3, and DENV4 (D1+3+4), S16803, QML-22, BSA, or un-depleted (None). Homotypic neutralization following each depletion condition was evaluated, including from NHPs infected with S16803 against S16803 (**A**) and NHPs infected with QML-22 against QML-22 (**B**). Heterotypic neutralization following each depletion was evaluated from NHPs infected with S16803 against QML-22 (**C**) and NHPs infected with QML-22 against S16803 (**D**). The values above the bars indicate the FRNT_50_ fold-change relative to the original antiserum sample (depletion condition “None”), which did not go through the depletion process. Arrows indicate meaningful (≥4-fold) reductions in FRNT_50_ titer. Error bars show 95% confidence intervals from technical duplicates. FRNT_50_ titers less than 80 were set to 40 (dashed line) on the graph.

We then evaluated how antisera depleted with each antigen neutralized the heterologous DENV2 strain. For S16803-inculated antisera, depletion with either S16803 or QML-22 substantially reduced neutralization of QML-22 (**Fig. 5C**). Similarly, QML-22-inoculated antisera lost neutralization of 16803 after depletion with either S16803 or QML-22 (**Fig. 5D**). The finding that depletion with either antigen could substantially reduce heterotypic DENV2 neutralization indicated that the strains shared one or more neutralizing determinants.

### The distinct residues in QML-22 are under relaxed selective pressure across serotypes

Given our finding that the 3F9 epitope was divergent in QML-22, we tested whether the 3F9 epitope was under differential selective pressure in the DENV E protein compared to other potently neutralizing epitopes. We first estimated site-specific non-synonymous to synonymous nucleotide substitution ratios (dN/dS) [36, 37]. Most sites across E were under negative selection, with neutral dN/dS ratios observed for 9 positions in DENV1, 16 in DENV2, 14 in DENV3, and 6 in DENV4 (**Fig. 6A**). DENV2 was under the strongest negative selective pressure, with a median dN/dS ratio of 0.032, and this differed significantly from DENV1 (0.049) and DENV4 (0.045) but not DENV3 (0.037) (Wilcoxon rank sum test with a Bonferroni correction, p-value <0.05). We then grouped site-specific dN/dS ratios by epitopes for 3F9 and other potently neutralizing antibodies (**Table S3**). For DENV2, the dN/dS distribution for residues in the 3F9 epitope was significantly greater than the EDE antibodies and the overall distribution of dN/dS for E (Wilcoxon rank sum test, p-value < 0.05) (**Fig. 6B**). Although the 3F9 epitope was not significantly different from additional epitopes in DENV2, the median was the highest of all epitopes, closely followed by the epitope for 5H2, a DENV4 type-specific antibody that targets many of the same E sites as 3F9. We also estimated the dN/dS ratios for the sites within these epitopes mapped to other serotypes and found the 3F9 and 5H2 epitopes were also under weaker negative selective pressure in DENV1 relative to EDE antibodies and 2D22.

**Figure 6.**
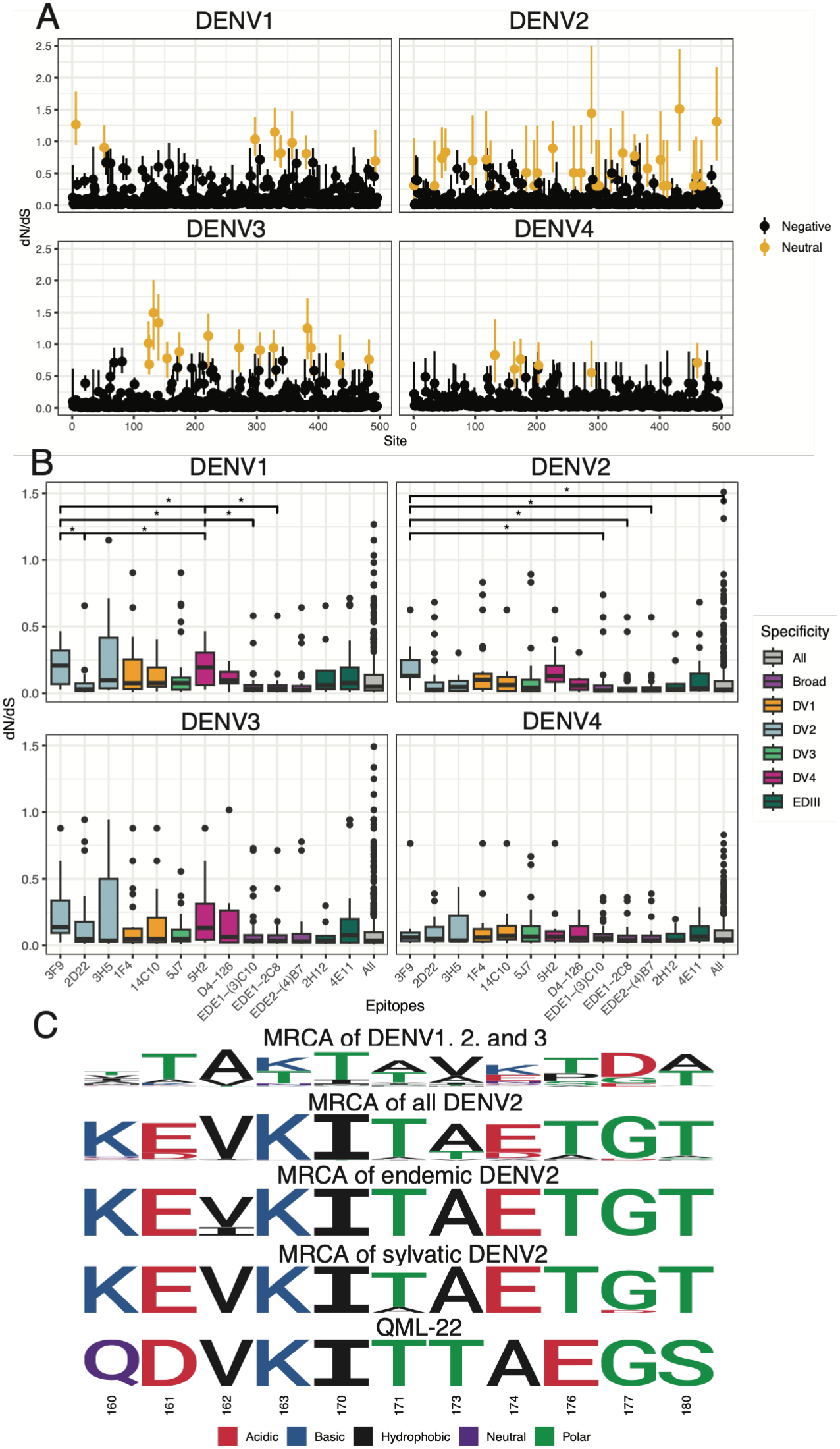
Evolutionary analysis of the 3F9 epitope in relation to other potent neutralizing epitopes. **(A)** DN/dS estimates per residue for each of the four DENV E proteins. Black denotes residues evolving under significant negative selection (with vertical bars indicating 95% High Density Region < 1.0); gold denotes sites with dN/dS with 95% High Density Region overlapping 1.0 (neutral). (**B**) The distribution of dN/dS values for each serotype for sites (y-axis) within well-characterized epitopes (x-axis) for n=824 sequences. Horizontal bars show medians, boxes indicate the interquartile range, and whiskers extend to 1.5 times the interquartile range. Distributions are colored by whether the epitope is serotype-specific or cross-reactive. “All” evaluates dN/dS across all sites in the E protein of that serotype. Differences across epitopes for each serotype were first evaluated by Kruskal-Wallis test, followed by group comparisons evaluated by Wilcoxon rank-sum test with a Bonferroni correction. * p<0.05; ** p<0.01; *** p<0.001; **** p<0.0001. (**C**) A sequence logo plot of the confidence of sequence identity of the predicted ancestral amino acids at sites within the DENV E 3F9 epitope. Bayesian Ancestral Sequence Reconstruction was performed on the ancestor of endemic, sylvatic, and all DENV2 E sequences as well as the common ancestor of DENV1, DENV2, and DENV3. Residues are colored by their biochemical class.

Given selection in DENV2 was relaxed at the 3F9 epitope, we evaluated if QML-22 retained ancestral residues in this region using Bayesian Ancestral Sequence Reconstruction. However, at positions 160, 161, 173, and 174 within the 3F9 epitope, QML-22 diverged from the ancestral population of both DENV2 and the common ancestor of DENV1, DENV2, and DENV3 in a manner not replicated by other lineages of DENV2, either endemic or sylvatic (**Fig. 6C**).

## Discussion

The dengue field remains uncertain how to classify strains of DENV that fall between the serotypes on phylogenetic trees [6]. All were isolated from patients infected on the island of Borneo and most are likely due to spillover from an unknown host reservoir. Here, we show one of these strains, QML-22, shares many antigenic features with other DENV2 strains, including recognition by potently neutralizing cross-reactive and some DENV2-specific antibodies. However, QML-22 also has unique amino acids within domain I of the E protein, which we show is a hotspot for evolution. These differences in QML-22 result in loss of neutralization by a potent DENV2 monoclonal antibody 3F9 and affect the immune response induced by QML-22. This study thus provides insight into the range of antigenic properties of DENV and the evolutionary pressures acting on different regions of the DENV E protein, both within and between serotypes.

Our findings suggest QML-22 is not only a phylogenetic intermediate between a genotype and serotype, but also an antigenic intermediate. One hypothesis is that EDE and many serotype-specific epitopes in domain III must be mostly conserved to maintain structural functions like virion formation and membrane fusion [38, 39], whereas the region of E domain I including the 3F9 epitope is both immunodominant and tolerant to mutations, with adaptation at this site enabling escape from preexisting immunity. Consistent with this hypothesis, many of the potently neutralizing type-specific antibodies identified to date against DENV1 (1F4 and 14C10) [40, 41], DENV2 (3F9) [31], DENV3 [42], and DENV4 (5H2) [43] are centered on this region of domain I. Further, this region defines differences between serotypes: changing amino acids 157, along with 126, can fully eliminate neutralization by antisera raised to DENV1 [44] and DENV3 naturally has deletions at positions 156 and 157 compared to other serotypes. Alternatively, evolution in this region of domain I may directly support adaptation to the vector and human by improving replicative fitness. In serial passage experiments, amino acids 150-160 in E domain I were under particularly strong positive selection in mosquito cells, due in part to modifying the N153 glycan [45]. In humans, a rare DENV2 lineage containing 160Q, the amino acid found in QML-22, resulted in higher viremia during secondary infection despite the 160Q virus being more susceptible to neutralization by heterotypic antisera in vitro [46].

From the initial antigenic data showing QML-22 was particularly well neutralized by heterotypic antisera, we hypothesized that QML-22 might be a ‘living fossil’ that had retained ancestral DENV features. However, our genetic and ancestral reconstruction analyses instead suggest it is more likely that QML-22 evolved in an immunological environment distinct from other DENV2 populations. QML-22 most likely circulates, primarily, in a sylvatic cycle, involving yet to be identified hosts, and, potentially, with slowed evolution due to periods of stasis in sylvatic mosquito eggs following transovarial transmission of virus [47]. This is supported by the original evolutionary analyses of QML-22, which found sylvatic DENV2 accumulated mutations at a lower rate than endemic DENV2 strains, and that QML-22 fell closer to endemic evolutionary rates, suggesting it may have a history of both endemic and sylvatic circulation [29]. Sylvatic strains more closely related to endemic strains have been isolated from NHPs in peninsular Malaysia [48], and spillover of sylvatic DENV2 into humans has been observed in Borneo [49]. Twenty eight per cent of Orangutans in the wild in Borneo were found to have anti-DENV2 antibodies, a similar proportion to that in the local human population (37%) [50], but the serological assays employed could not have distinguished between infection with endemic or sylvatic strains of virus. In Senegal, persistent transmission of DENV2 among NHPs [51], results in occasional outbreaks in humans, most recently in 2020 [52]. Given that two of the four divergent DENV strains from Borneo were isolated from international travelers, not local residents, it is likely that the burden of spillover of divergent DENV strains like QML-22 is underreported. Rapid deforestation to provide land for agriculture increases opportunities for humans to interact with bridge vectors, such as *Aedes niveus* and *Aedes albopictus* [53]. Development in rural eastern Borneo, such as Ibu Kota Nusantara, the new capital of Indonesia, may further increase opportunities for spillover of sylvatic and highly divergent DENV strains as well as other zoonotic pathogens [54].

Our study has several limitations. Given the laborious nature of depletion experiments, only n=2 NHPs were depleted per infecting antigen. Further, our analyses focused on the E protein, given the importance of E to viral antigenic properties. However, other viral proteins may also differ in important ways between QML-22 and endemic DENV2 strains. Additionally, the evolutionary analyses focused on known quaternary anti-E epitopes; there are likely other relevant neutralizing epitopes not studied here. Finally, we only had QML-22 available to us and could not compare it to other divergent strains. Future studies should compare across divergent strains to better understand their shared features.

The pandemic potential of divergent strains like QML-22, Brun2014, DKE-121, and UDS37/9 is uncertain but most likely low. Available data suggests these viruses share many serotype-specific and conserved cross-reactive epitopes, and thus endemic DENV infection is likely to confer protection. Yet, the amino acid differences within E domain I in QML-22 are also sites of endemic genotype-level variation, and prior studies have shown vaccination is often more genotype-specific, potentially focusing on more divergent epitopes like those in domain I [14]. Given that antigenic escape from specific epitopes is one of the fitness effects that likely contributes to successful DENV lineage introductions [17], the antigenic difference of known and unknown divergent DENV strains is potentially relevant. Further study of these and other divergent strains may provide insight into the antigenic properties of the four DENV serotypes, including sites of likely future evolution, and the factors driving their continued antigenic diversification.

## Materials and Methods

### QML-22 virus and phylogenetic comparisons

The QML-22 strain was sequenced as described previously (KX274130) and also sequenced by a separate laboratory (KY923048) [29, 55]. Sequence analysis was performed on QML-22, DKE-121 (MZ215848), Brun2014 (KR919820), and UDS37-09 (EPI_ISL_20063516) [27], and globally representative endemic and sylvatic strains [6, 56], including those that have been previously antigenically characterized [17, 18]. Sequence alignments were made in MAFFT, and phylogenetic trees were made using IQ-Tree with model selection to identify GTR+F+R7 as the best fit. Tree plots were generated with the ggtree package in R. Percent dissimilarity from QML-22 was estimated relative to all global DENV2 strains. Differences were plotted on the E protein structure from Protein Data Bank #: 6ZQU.

### Antigenic comparison of QML-22 to a global DENV1-4 panel

Antigenic characterization of QML-22 in relation to globally representative [17] and additional contemporary strains [18] of DENV1-4 was performed by testing how QML-22 was neutralized by 20 reference NHP antisera using immunofocus reduction neutralization test (FRNT). The reference antisera were generated by inoculating each African green monkey (*Chlorocebus sabaeus*) subcutaneously with 10^5^ immunofocus forming units (FFU) of a different strain from the globally representative DENV1-4 panel [17]. Sera or plasma were collected 90 days after inoculation. Two additional African green monkeys were inoculated subcutaneously with 10^5^ FFU of low-passage QML-22. Due to low initial homologous neutralizing antibody titers, at seventy days post-inoculation, both animals were given a second dose of 10^5^ FFUs. Serum collected at day 30 post-boost (day 100 post-initial inoculation) was used to test neutralization of QML-22 and other DENV1-4 strains. Animals were infected on protocols approved by the NIAID Animal Care and Use Committee. We followed the National Institutes of Health guidelines for the humane treatment of laboratory animals.

The FRNT for the antigenic cartography analyses was performed as previously described on C6/36 mosquito cells (CRL-1660, ATCC) propagated at 32°C [17, 18]. Two-fold dilutions of antisera were mixed with constant amount of virus in diluent (OptiMEM, 2% FBS, 50 μg/mL gentamicin, and 0.5% albumin) and incubated at 37°C for 1.5 hours. Virus-antibody mixtures were added in technical duplicates to cells and incubated at 32°C for 1 hour. After incubation, an overlay of 1% methylcellulose (Sigma-Aldrich) in OptiMEM GlutaMAX (2% FBS, 2.5 μg/mL amphotericin B, 20 μg/mL ciprofloxicin) was added in each well. The plates were incubated at 37°C for 3 days. Cells were washed with PBS, fixed 80% methanol, and immunostained using anti-DENV antibodies 2H2 and 4G2, Peroxidase-labeled goat anti-mouse IgG (Kirkegaard & Perry Laboratories, KPL, Gaithersburg, MD), and stained with TrueBlue Peroxidase substrate (KPL). Plates were imaged with the ImmunoSpot Analyzer (CTL, Shaker Heights, OH) and immunofoci counted using the Viridot package in R [17, 18, 57]. FRNT_50_ titers were estimated using the dcr package in R.

The viruses included on prior antigenic maps are as described previously [17, 18]. Antigenic maps using this previously collected data were made to incorporate QML-22. Maps were made using a modified multidimensional scaling method, which treats neutralization titers as measures of antigenic distance between each serum and virus [58]. Antigenic distance *Dij* between virus *i* and antiserum *j* was defined by subtracting the neutralization titer for *i* and *j*, *Nij*, from the titer for the virus best neutralized by that antiserum, defined as *bj*: *Dij* = log_2_(*bj*)-log_2_(*Nij*). Antigenic map positions were identified by finding the Euclidean distance *dij* that best fit the measured antigenic distance *Dij*. One unit of map antigenic distance was directly proportional to a two-fold difference in FRNT50 titers. To create the antigenic map, the error function E=∑*ij*e(*Dij*,*dij*) was minimized from random starting coordinates using L-BFGS gradient-based optimization. The minimum error map was presented from 10,000 independent optimizations. Maps were made using the Racmacs package in R.

### Breathing and maturation experiments

Breathing experiments were designed as described by others [23]. Monoclonal antibodies C8, C10, and 4G2 were produced with Curia, and E60 was kindly provided by Dr. Theodore Pierson at the National Institutes of Health. QML-22 was compared to DENV2 Tonga (MW946475) and DENV2 Cambodia (KT452795), which captured the range of DENV2 antigenic variation observed on antigenic map. Viruses were propagated in C6/36 cell cultures to produce standard virus stocks, as described previously [17, 18, 57]. Mature virus stocks were grown in Vero-81 cells (originally CCL-81, ATCC, Manassas, VA) engineered to overexpressing furin [59]. Vero cells were propagated at 37°C and 5% CO_2_ in Opti-Pro serum-free medium (GIBCO, Life Technologies, Carlsbad, CA) supplemented with 200uM glutamine (Sigma-Aldrich, Saint Louis, MO), 100 U/mL penicillin and 100 mg/mL streptomycin (GIBCO). To evaluate virion breathing, constant amount of virus and serial dilutions (starting at 10 µg/mL) of monoclonal antibodies were incubated at 37°C for 1.5 or 4-hours. Virus-antibody mixtures were then added in technical duplicates to C6/36 cells seeded in 96-well plates, following the neutralization assay procedure described above [17, 18, 57]. In addition, the mature DENV2 strains were tested against the NHP antisera on C6/36 cells using the 1.5 serum-virus incubation period, and the mature and standard stocks were tested using an FRNT with conventional Vero-81 cells as the substrate (CCL-81, ATCC) propagated and maintained at 37°C as described above, but otherwise following the same FRNT protocol as used for C6/36 cells [17, 18, 57].

### Serum depletion experiments

Neutralizing activity of depleted antisera (described below) and the DENV2 monoclonal antibodies 4E11, 2D22, 3H5, and 3F9 were tested by FRNT as described above, with the following differences. Vero-81 cells were propagated at 37°C and 5% CO_2_ in Dulbecco’s Minimal Essential Medium (DMEM) (GIBCO), supplemented with 10% fetal bovine serum (FBS) (Sigma-Aldrich, Saint Louis, MO), 100 U/mL penicillin and 100 mg/mL streptomycin (GIBCO). Four-fold dilutions of antisera or antibodies were mixed with constant amount of virus and incubated at 37°C for 1 hour. Virus-antibody mixtures were added to cells in technical duplicates and incubated at 37°C for 1 hour. After incubation, an overlay of 0.8% methylcellulose (Sigma-Aldrich) in DMEM (supplemented with 1% FBS, 100 U/mL penicillin and 100 mg/mL streptomycin) was added in each well. The plates were incubated at 37°C for 2 to 3 days depending on the virus phenotype. Cells were washed with PBS and fixed with acetone and methanol (1:1) solution and immunostained as described above. Data were analyzed by nonlinear regression to determine 50% neutralization titers in GraphPad Prism 8. Monoclonal antibody 3H5 was obtained from Envigo, 4E11 from GenScript, and 2D22 and 3F9 kindly provided by Dr. Aravinda de Silva at the University of North Carolina School of Medicine.

DENV2 S16803 rhesus macaques (*Macaca mulatta*) infections were conducted under an IACUC-approved animal use protocol in an AAALAC International - accredited facility with a Public Health Services Animal Welfare Assurance and in compliance with the Animal Welfare Act and other federal statutes and regulations relating to laboratory animals.

To produce high titer stocks of antigen, viruses were propagated in C6/36 cells at 28°C and 5% CO_2_ in DMEM supplemented with 10% FBS, 100U/mL penicillin-streptomycin and 1% tryptose phosphate broth (TPB) (Sigma-Aldrich, Saint Louis, MO). Supernatants were concentrated through AMICON filters (100 KDa MWCO, Millipore) and then purified on a sucrose gradient. Antiera from rhesus macaques immunized with DENV2 S16803 [35] or QML-22 (see above) were depleted of antibodies as previously described [35]. Briefly, 0.2 mL of polystyrene cross-linked magnetic beads (Spherotech) were coated overnight with constant rotating at 4°C with 150 µg of purified QML-22, S16803, or a combination of DENV1 (West Pac 74), DENV3 (CH53489) and DENV4 (TVP-360), or BSA (control). After incubation, antigen:bead complexes were blocked with BSA (10 mg/mL) in PBS for 2 hours at room temperature and washed twice in PBS by magnetic separation. Heat-inactivated immune antisera at 1:20 dilution in PBS were then incubated for 1 hour at 37°C with the antigen:bead complexes with constant rotating. Depletions were performed by transferring the antisera to new antigen:bead complexes. Two independent depletion experiments were conducted and yielded similar results. The results shown here are following four rounds of depletion.

Antibody depletion was confirmed by ELISA [35]. Briefly, 96-well ELISA plates were coated overnight at 4°C with 50µL (2µg/well) of purified DENV antigen or BSA, blocked for 1 hour at room temperature (RT) in 5% non-fat dry milk and 0.25% BSA in PBS-T solution and assayed with the depleted serum samples in technical duplicates starting at 1:200 dilution for 1 hour at RT. Secondary anti-human IgG-HRP (Sigma-Aldrich) antibody was added and incubated for 1 hour at RT. Then 3,3’,5,5’-tetramethylbenzidine (TMB) substrate (SeraCare) was added, incubated until color development, and the reaction stopped with addition of 1M sulfuric acid. Absorbance was measured at OD 450nm.

### Selective pressure analyses

An expanded alignment of DENV C, prM, and E sequences was downloading from NCBI by searching “DENV” on December 16th, 2025, grouped by serotype. CD-HIT with a similarity threshold of 98% was used to subset the collection of sequences, then used Clustal Omega for alignment. The resulting alignment was then manually curated before being trimmed to contain only C, prM, and E. Finally, we performed an additional round of CD-HIT at a 99% threshold followed by codon alignment in Aliview. We used IQ-Tree to generate a phylogenetic tree of the expanded alignment of 824 DENV C, prM, and E nucleotide sequences. We used ultrafast bootstrapping. All structures were visualized with ChimeraX version 1.9 [60]. For dN/dS analysis, we implemented a BEAST analysis [36, 37] on each serotype independently. We generated an XML with an HKY substitution model and exponential growth set as a tree prior. We employed Renaissance counting to reconstruct site dN/dS change counts with an MCMC chain of length 100,000,000 and logging every 10,000 states. The BEAST run for each serotype included 3 replicates, and the dN/dS values were averaged per site. For epitope analysis, we used residues listed in the DENV-Ab database [61]. Plots were generated with ggplot2.

## Acknowledgments

We thank the researchers who deposited the sequences used in these analyses in both GenBank and GISAID. We thank Dr. Theodore Pierson of the National Institutes of Health for providing monoclonal antibody E60 and Dr. Aravinda de Silva at the University of North Carolina School of Medicine for providing monoclonal antibodies 2D22 and 3F9. This research was supported by the Intramural Research Program of the National Institutes of Health (NIH), the National Institute of Environmental Health Sciences Health and Extreme Weather Intramural Research Program, NIH grants R01AI114703-01, AI106695-12, P01AI106695, and U19AI181960, and the Military Infectious Disease Research Program. The contributions of the NIH authors were made as part of their official duties as NIH federal employees, are in compliance with agency policy requirements, and are considered Works of the United States Government. However, the findings and conclusions presented in this paper are those of the authors and do not necessarily reflect the views of the NIH or the U.S. Department of Health and Human Services. Material has been reviewed by the Walter Reed Army Institute of Research. There is no objection to its presentation and/or publication. The opinions or assertions contained herein are the private views of the author, and are not to be construed as official, or as reflecting true views of the Department of the Army or the Department of Defense.

## Data availability

All data are presented in the manuscript and supplementary materials. The raw data and code used for making the figures will be deposited in Zenodo (10.5281/zenodo.20631407) and made available at the time of publication. Correspondence and requests for materials should be directed to L.C.K. and G.D.G.

