## Supplementary information for "Evolutionarily distinct dengue virus type 2 with a unique type-specific neutralization determinant"

### Supporting Information

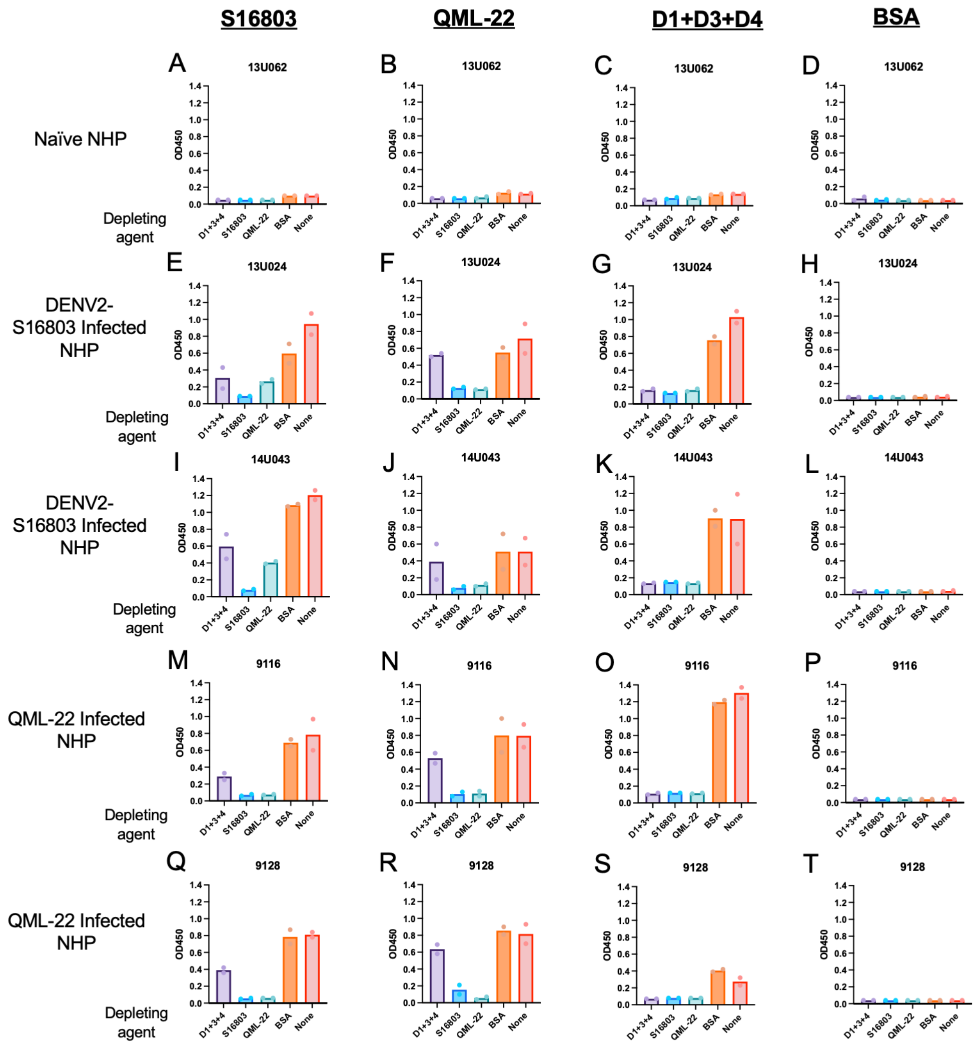


**Figure S1. ELISAs with antibody-depleted NHP antisera.** Individual data points (dots) and average (bars) optical densities at 450nm (OD450) for technical duplicates of each depleted serum and ELISA antigen combination are shown. The antigens used for the ELISA were purified virions of DENV2 S16803 (panels A, E, I, M, and Q), purified virions of DENV2 QML-22 (panels B, F, J, N, and R), a combination of purified virions of DENV1 West Pac 74, DENV3 CH53489, and DENV-4 TVP-360 (panels C, G, K, O, and S), and bovine serum albumin (BSA)

(panels D, H, L, P, and T). The serum antibody depletion conditions are indicated below each bar. Depletion condition “None” is original antisera that did not go through the depletion process. The NHP serum samples are listed above each graph. Animal 13U062 was flavivirus immunologically naïve (panels A-D), animals 13U024 and 14U043 were previously infected with DENV2 S16803 (panels E-L), and animals 9116 and 9128 were previously infected with QML-22 (panels M-T).

**Table S1. Alignment showing amino acids with a higher percentage of difference between QML-22 and other globally representative DENV2 strains (n=843 sequences).**

| E residue | AA in QML-22 | # strains same as QML-22 | % Dissimilarity from QML-22 | Most common AA in DENV2 | % of strains with this AA |
|---|---|---|---|---|---|
| 4 | V | 0 | 100.0 | I | 100.0 |
| 7 | A | 0 | 100.0 | S | 100.0 |
| 46 | T | 5 | 99.4 | I | 98.9 |
| 52 | L | 1 | 99.9 | Q | 82.4 |
| 83 | P | 0 | 100.0 | K | 60.7 |
| 91 | I | 81 | 90.3 | V | 90.1 |
| 93 | R | 20 | 97.6 | K | 97.5 |
| 96 | L | 0 | 100.0 | M | 93.7 |
| 124 | Y | 1 | 99.9 | N | 97.4 |
| 139 | V | 7 | 99.2 | I | 99.0 |
| 140 | A | 0 | 100.0 | V | 99.8 |
| 141 | V | 537 | 36.1 | V | 63.9 |
| 149 | H | 637 | 24.3 | H | 75.7 |
| 150 | S | 1 | 99.9 | A | 99.4 |
| 160 | Q | 1 | 99.9 | K | 92.9 |
| 161 | D | 0 | 100.0 | E | 100.0 |
| 162 | V | 41 | 95.1 | I | 95.0 |
| 164 | I | 151 | 82.0 | V | 82.0 |
| 173 | T | 6 | 99.3 | A | 99.0 |
| 174 | A | 1 | 99.9 | E | 99.9 |
| 176 | E | 0 | 100.0 | T | 99.3 |
| 180 | S | 0 | 100.0 | T | 99.8 |
| 181 | I | 27 | 96.8 | V | 95.7 |
| 182 | S | 0 | 100.0 | T | 99.9 |
| 203 | D | 128 | 84.8 | N | 81.9 |
| 224 | S | 0 | 100.0 | A | 99.9 |
| 226 | T | 478 | 43.1 | T | 56.9 |
| 228 | G | 566 | 32.5 | G | 67.5 |
| 232 | V | 3 | 99.6 | I | 99.6 |
| 262 | S | 0 | 100.0 | T | 99.8 |
| 273 | T | 0 | 100.0 | S | 99.9 |
| 293 | E | 0 | 100.0 | Q | 99.9 |
| 329 | E | 9 | 98.9 | D | 98.8 |

| | | | | | |
|---|---|---|---|---|---|
| 345 | K | 23 | 97.3 | R | 97.3 |
| 346 | H | 330 | 60.7 | Y | 60.5 |
| 358 | I | 4 | 99.5 | V | 99.0 |
| 363 | K | 0 | 100.0 | S | 95.0 |
| 365 | T | 0 | 100.0 | V | 96.9 |
| 379 | L | 0 | 100.0 | I | 97.0 |
| 389 | M | 0 | 100.0 | L | 100.0 |
| 430 | N | 0 | 100.0 | T | 100.0 |
| 443 | V | 3 | 99.6 | I | 99.6 |
| 445 | N | 0 | 100.0 | G | 99.9 |
| 462 | I | 620 | 26.4 | I | 73.6 |
| 475 | M | 0 | 100.0 | L | 100.0 |
| 477 | I | 0 | 100.0 | V | 100.0 |
| 478 | T | 25 | 97.0 | S | 97.0 |
| 481 | M | 0 | 100.0 | L | 99.6 |
| 484 | L | 0 | 100.0 | I | 64.9 |
| 492 | V | 14 | 98.3 | M | 98.0 |

**Table S2. Ability of antisera from naïve, QML-22-infected, and S16803-infected NHP antisera before and after antibody-depletion to neutralize QML-22, S16803, and DENV1, DENV3, and DENV4.**

| NHP (DENV2 used for infection) | Virus | FRNT$_{50}$ titer after depletion with: | | | | |
|---|---|---|---|---|---|---|
| | | D1+3+4 | S16803 | QML-22 | BSA | None |
| 13U062 (none) | DENV2 S16803 | <80 | <80 | <80 | <80 | <80 |
| 13U024 (S16803) | | 13858 | 1332 | 11855 | 13810 | 24301 |
| 14U043 (S16803) | | 7722 | 348 | 7052 | 9990 | 10779 |
| 9116 (QML-22) | | 190 | <80 | <80 | 330 | 316 |
| 9128 (QML-22) | | 612 | <80 | <80 | 5038 | 1865 |
| 13U062 (none) | DENV2 QML-22 | <80 | <80 | <80 | <80 | <80 |
| 13U024 (S16803) | | 1425 | 176 | 297 | 2205 | 2220 |
| 14U043 (S16803) | | 934 | <80 | 129 | 2089 | 2546 |
| 9116 (QML-22) | | 946 | 499 | 249 | 986 | 1205 |
| 9128 (QML-22) | | 474 | 248 | 110 | 822 | 752 |
| 13U062 (none) | DENV1 WP74 | <80 | <80 | <80 | <80 | <80 |
| 13U024 (S16803) | | <80 | <80 | <80 | 222 | 113 |
| 14U043 (S16803) | | <80 | <80 | <80 | <80 | <80 |
| 9116 (QML-22) | | <80 | <80 | <80 | <80 | <80 |
| 9128 (QML-22) | | <80 | <80 | <80 | <80 | <80 |
| 13U062 (none) | DENV3 CH53489 | <80 | <80 | <80 | <80 | <80 |
| 13U024 (S16803) | | <80 | <80 | <80 | <80 | <80 |
| 14U043 (S16803) | | <80 | <80 | <80 | <80 | <80 |
| 9116 (QML-22) | | <80 | <80 | <80 | <80 | <80 |
| 9128 (QML-22) | | <80 | <80 | <80 | <80 | <80 |
| 13U062 (none) | DENV4 TVP-360 | <80 | <80 | <80 | <80 | <80 |
| 13U024 (S16803) | | <80 | <80 | <80 | <80 | 125 |
| 14U043 (S16803) | | <80 | <80 | 102 | <80 | <80 |
| 9116 (QML-22) | | <80 | <80 | <80 | <80 | <80 |
| 9128 (QML-22) | | <80 | <80 | <80 | <80 | <80 |

**Table S3. Previously mapped contact or escape residues for monoclonal antibodies plotted in Fig. 6B, derived from the DENV-Ab database.**

| Specificity | Antibody name | Residue in the E protein in contact with antibody |
|---|---|---|
| DENV2 | 3F9 | 160, 161, 162, 163, 170, 171, 173, 174, 176, 177, 180 |
| DENV2 | 2D22 | 2, 66-74, 99, 101-105, 113, 148-149, 151-155, 246-248, 307, 309, 310, 325, 327, 360-364 |
| DENV2 | 3H5 | 301, 305, 383, 384, 327, 329, 332 |
| DENV1 | 1F4 | 46-49, 51-52, 136, 138, 155-157, 160-163, 165, 170-174, 176, 177, 273-276 |
| DENV1 | 14C10 | 51, 53, 134, 136, 160-163, 167, 168, 171-174, 273-276, 278 |
| DENV3 | 5J7 | 50-55, 58, 73, 74, 101, 106, 123, 126, 128, 130, 131, 133, 134, 148, 196, 198, 200, 201, 223, 224, 227, 274, 276, 307-309 |
| DENV4 | 5H2 | 160-163, 168, 170-174, 176, 177, 179, 180, 291, 293 |
| DENV4 | D4-126 | 51, 53, 124, 135, 200, 234 |
| Broad | EDE1-(3)C10 | 2, 27-29, 44-47, 67-74, 77, 81-84, 97-100, 101-107, 113, 115, 246-249, 273-274, 309-310, 323, 325, 362 |
| Broad | EDE1-2C8 | 67-74, 77, 83-84, 98-100, 102-107, 113, 158, 245-248, 273-274, 310-311 |
| Broad | EDE2-(4)B7 | 67-74, 98-100, 102-105, 113, 152-156, 157, 245-249 |
| EDIII complex | 2H12 | 310, 313-317, 319, 321, 323, 352-354, 363, 364, 366, 368, 370 |
| EDIII complex | 4E11 | 305-312, 323, 325, 327, 360-364, 385-391 |
